# Physics-aware resolution enhancement of soft X-ray tomography with measurement-supervised learning

**DOI:** 10.64898/2026.06.21.730079

**Authors:** ShaoSen Chueh, Eoghan Gallagher, Charlotte de Ceuninck van Capelle, Leo Luo, Takashi Ishikawa, Christopher Evans, Nicola Fletcher, Mary Lopez-Perez, David Rogers, Stephen O’Connor, Conal McIntyre, Martina Donnellan, Paul Sheridan, Jeremy C. Simpson, Sergey Kapishnikov

## Abstract

Soft X-ray tomography (SXT) is an emerging modality for whole-cell 3D imaging in near-native states. However, the effective spatial resolution is limited by optical artifacts characterized by the point spread function (PSF). Standard reconstruction methods force a compromise between structural sharpness and noise, failing to fully resolve these depth-dependent artifacts. By embedding experimentally measured, depth-variant PSFs into a differentiable forward model, we demonstrate a physics-aware computational optimization that bypasses these limitations to recover high-frequency cellular ultrastructure. The structural fidelity was validated using split-tilt Fourier ring correlation (FRC), alongside an experimental bead phantom tomogram, providing supporting evidence that the recovered high-frequency features reflect genuine specimen structure rather than fabricated artifacts. Our method effectively increases FRC spatial resolution and recovers cellular ultrastructure. Furthermore, under sparse-angular subsampling, the framework maintained spatial resolution using half the projection angles, a computational proxy pointing toward the potential for reduced radiation exposure in future acquisitions. This hardware-free, computational approach offers a route toward mitigating the optical and dosimetric constraints that currently limit nanoscale soft X-ray tomography.

## Introduction

Soft X-ray tomography (SXT) is a label-free imaging modality capable of capturing volumetric features of intact cells in their near-native states. By resolving three-dimensional cellular ultrastructure, SXT enables visualization of the entire cellular volume and its internal spatial organization, providing structural data for biological and biomedical research(Harkiolaki et al., 2018; Kapishnikov et al., 2021; Nahas et al., 2022; Huang et al., 2024; McDermott et al., 2009; Loconte et al., 2022; Guo & Larabell, 2019; Schneider et al., 2010).

However, effective SXT resolution is affected by multiple factors, including beam monochromaticity, partial coherence, mechanical stability, sample vitrification, and strict radiation dose limits, where gaining a two-fold improvement in resolution requires a sixteen-fold increase in photon exposure. Among the factors to achieve nanometer-scale resolution, objective lenses with high numerical apertures (NA) are required. However, this introduces a physical trade-off: the depth of field (DOF) decreases quadratically with increasing NA. This physical phenomenon violates the standard parallel-projection assumptions used in conventional tomography(Radermacher, 1992; Radon, 1986; Gilbert, 1972), introducing depth-variant defocus blur on the biological specimens with thickness exceeding the restricted DOF(Weiß et al., 2000; Hagen et al., 2012; Duke et al., 2014; Chen et al., 2016; Otón et al., 2017; Oton et al., 2012; Ekman et al., 2018). Cellular structures exhibit gradual optical degradation with increasing distance from the focal plane, which compromises the integrity of the 3D reconstruction. Because these optical artifacts are generated during imaging, the quality of 2D projections (tilt series) is degraded from optimal resolution.

Resolving this complicated degradation remains a great computational and engineering challenge. Conventional tomographic reconstruction algorithms, which rely on depth-invariant ray approximations, often yield distortions and loss of resolution in these defocused regions(Radermacher, 1992; Radon, 1986; Gilbert, 1972). While recent hardware-driven (or acquisition-driven) strategies, such as half-acquisition geometry combined with broadband illumination, have expanded the lateral field of view for larger cells, residual defocus artifacts may still manifest at the specimen periphery(Ekman et al., 2018). Furthermore, this technique requires full-rotation stages and capillary substrates, making its application incompatible with flat-grid cryo-SXT workflows that are constrained by limited tilt angles. Similarly, through-focus multiplexing techniques such as XTEND(Otón et al., 2017) can computationally extend the DOF by acquiring focal series at each tilt angle; however, this technique requires data collection at different focal positions.

On the other hand, purely computational strategies have struggled to resolve this depth-dependent blur. Simplified approaches, such as applying 2D depth-independent Wiener deconvolution to projection images(Otón et al., 2016), fail to model the true 3D depth-dependent nature of the degradation. Furthermore, incorporating the 3D depth-variant point spread function (PSF) into a linear projection matrix makes the inverse problem highly unstable when realistic measurement noise is present(Ekman et al., 2018). While classical linear inversion methods can partially reduce this instability through iterative regularization, such as early stopping, they are limited by a mathematical trade-off. Therefore, solving this ill-posed inverse problem while maintaining sharp biological features remains challenging.

While deep learning has recently achieved tremendous success in microscopy image processing and analysis(Falk et al., 2019; Wang et al., 2019; Rong Chen et al., 2023; Archit et al., 2025; Ouyang et al., 2018), applying purely data-driven models to highly ill-posed inverse problems poses risks of generating unfaithful features. Consequently, in computational imaging, the paradigm has transitioned towards physics-aware frameworks. By integrating governing physical rules into the neural networks or the training schemes, researchers can constrain the model output with physics rules and prevent hallucinations(Ratner et al., 2021; Page & Favaros, 2020; Luo et al., 2025; Yao et al., 2022; Li et al., 2025; Raissi et al., 2019; Chen et al., 2025; Ni Chen et al., 2023).

To address these optical and dosimetric constraints, the objective of this work is to apply an instance-specific optimization framework inspired by deep image prior(Ulyanov et al., 2020) that mitigates the SXT point spread function without requiring hardware or image acquisition modifications. By employing a physics-aware optimization strategy constrained by a differentiable SXT forward model that is constructed directly from an experimentally measured, depth-variant PSF, the hallucination risks inherent in purely data-driven super-resolution networks(Saharia et al., 2022) can be minimized. However, we acknowledge that this constraint does not eliminate the priors embedded in the network architecture, pretraining distribution, or initialization; rather, it provides an additional, physically grounded criterion against which the recovered structure can be evaluated.

In this work, we demonstrate that instance-specific approach consistently improves resolution across the specimens tested, spanning four distinct cell types. Furthermore, we show that, under retrospective sparse-angular subsampling, our optimization maintains spatial resolution using half the projection angles, a computational proxy pointing toward reduced radiation exposure, which could minimize biochemical degradation and mass loss in biological specimens(Gianoncelli et al., 2015; Shinohara & Ito, 1991). Ultimately, this framework aims to provide a computational solution to minimize radiation damage and maximize imaging throughput in nanoscale tomography.

## Materials and Methods

### SXT Image Formation Forward Model

In soft X-ray tomography (SXT), achieving nanometer-scale resolution necessitates an optical geometry that produces a shallow depth of field (DOF) relative to the specimen thickness. Because intact biological samples typically exceed this focal depth, structures outside the focal plane are projected onto the detector with depth-dependent blurring. To physically constrain our neural network, we implemented a forward model that maps the 3D linear absorption coefficients (LAC) of the cellular specimen, *μ*(*x,y,z*), to the 2D detector by incorporating this spatial degradation.

At each depth along the optical axis (*z*), the local absorption profile is spatially convolved with a 2D Point Spread Function (PSF), *h*_*z*_(*x,y*), dictated by its exact distance from the focal plane:

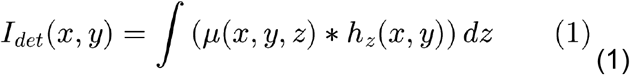

To accurately formulate *h*_*z*_ for the laboratory SXT-100 system, we measured the optical response of the microscope using a nanoscale Siemens star test pattern (Fig. 1a). The pattern was imaged at 9 discrete focal planes along the optical axis, spanning from − *μm* to +2.0*μm* in 0.5*μm* increments. However, the true, absolute focal plane (*z* = 0) is rigorously identified post-acquisition by selecting the plane that exhibits the highest peak spatial intensity along the optical axis.

**Figure 1.**
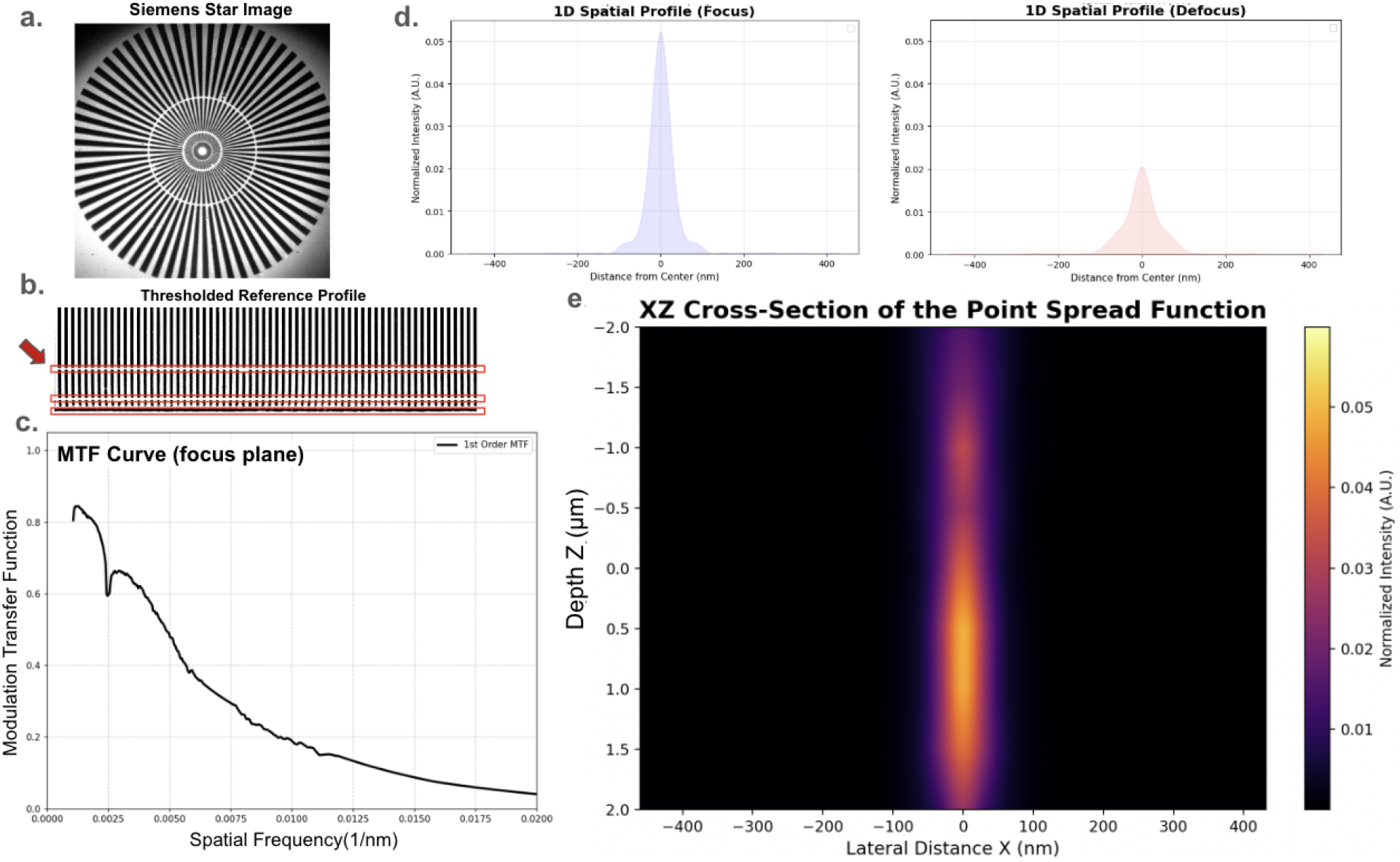
Experimental derivation of depth-variant Point Spread Functions (PSF). a) Original soft X-ray projection image of the nanoscale Siemens star test pattern. b) The thresholded binary reference mask profile, with red boxes highlighting the automatically detected support rings and unresolved inner radii that were excluded from the analysis via margin dilation. (c) The stabilized 1D MTF curve, mapped to true physical spatial frequency and interpolated across the physical pattern gaps. (d) 1D spatial intensity profiles comparing the Point Spread Function at the focal plane versus a defocus plane. (e) 2D XZ cross-section (depth profile) of the PSF, visualizing the continuous spatial blurring along the optical axis. (Note: The Z-axis labels represent relative distances from the initially estimated focus; the true focal plane is determined by the point of maximum intensity).

Following the methodology established by Otón et al.(Otón et al., 2015), these 9 images were processed to extract the Modulation Transfer Function (MTF). The circular Siemens star images were transformed into polar coordinates to linearly map spatial frequencies. To isolate the highest-quality signal, the polar-unwrapped images were cropped to retain the full angular width of the pattern while discarding the upper region containing dark artifacts introduced by the polar unwrapping process. To account for uneven illumination without blowing out bright edges, a localized contrast stretch was computed using the central region of the image and applied globally. A 1D Fast Fourier Transform (FFT) was then applied row-by-row to extract the magnitude of the 1st order (fundamental) harmonic from this isolated region.

Manufactured Siemens star patterns contain concentric support rings that interrupt the periodic spokes, and at extreme inner radii, optical blurring completely merges the adjacent spokes into an unresolved region devoid of periodic structural information. An automated gap-detection algorithm was implemented to identify these regions. To prevent noise spikes generated by the half-corrupted rows at the edges of these physical boundaries, a safety margin (3 pixel rows) was applied to the detected gaps. These excluded support rings and the fully unresolved central rows are illustrated as red exclusion zones on the ideal binary reference mask (Fig. 1b).

To calculate the MTF, the measured 1st order magnitude at each valid radius was normalized array-by-array against the 1st order magnitude extracted from the corresponding row of the ideal binary reference mask. Relying exclusively on the 1st-order harmonic prevents the SNR degradation associated with averaging higher-order harmonics. To transition from raw pixel radii to true physical spatial frequencies (*nm*^−1^), the radial data was scaled using the known geometric properties of the Siemens star (71 spokes) and the system’s pixel size. The resulting discrete MTF data was linearly interpolated across the excluded gap regions to yield a single, continuous, and physically accurate 1D MTF curve for each focal plane (Fig. 1c).

To transition from these frequency-domain contrast metrics to spatial blurring kernels, the 1D MTF profiles were converted into 2D PSFs(Otón et al., 2016). Assuming rotational symmetry for the objective lens, the 1D MTF data was extrapolated to 100% contrast at zero frequency and radially expanded to generate a 2D Apparent Transfer Function (ATF). An inverse 2D Fast Fourier Transform (iFFT) was applied to the ATF, yielding the spatial PSF (*h*_*z*_) for each measured depth. The resulting 1D spatial intensity profiles demonstrate a sharp, intense localization at the focal plane that distinctly broadens and flattens at defocused planes (Fig. 1d). Aggregating these normalized 1D profiles along the optical axis visualizes the characteristic spatial blurring natively produced by the objective lens (Fig. 1e).

These 9 experimentally derived PSFs function as depth-resolved spatial anchors in our forward model. During the forward projection of a 3D computational volume, the physical depth (*z*) of each voxel slice is calculated based on the system’s voxel size. For intermediate slices falling between the 9 measured planes, the algorithm applies linear interpolation between the two nearest 2D PSFs in the frequency domain. This establishes a continuous, physically constrained gradient of blurring kernels across the entire optical axis to simulate the out-of-focus degradation of the SXT-100.

Finally, to address the pulse-to-pulse energy jitter and Poisson-Gaussian statistics inherent to the laser-produced plasma (LPP) source, we integrated the Noise2Noise framework to extract the intrinsic noise distribution (*η*) directly from the soft X-ray tilt series(Lehtinen et al., 2018; Chueh et al., 2025). By combining the experimentally measured, interpolated PSF with data-driven noise modeling, the complete differentiable forward model is formulated as:

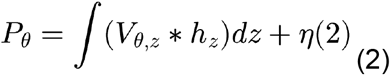

where *P*_*θ*_ is the simulated projection at tilt angle *θ, V*_*θ,z*_ is the rotated 3D volume slice at depth *z*, and *h*_*z*_ represents the experimentally measured, depth-variant PSF. This formulation ensures that the spatial degradation during neural network optimization is strictly governed by the measured optical physics of the instrument.

### Measurement-Supervised Network Optimization

With the differentiable forward model *F* defined in equation (2), we now integrate it into our complete network optimization pipeline (Fig. 2b).

**Figure 2.**
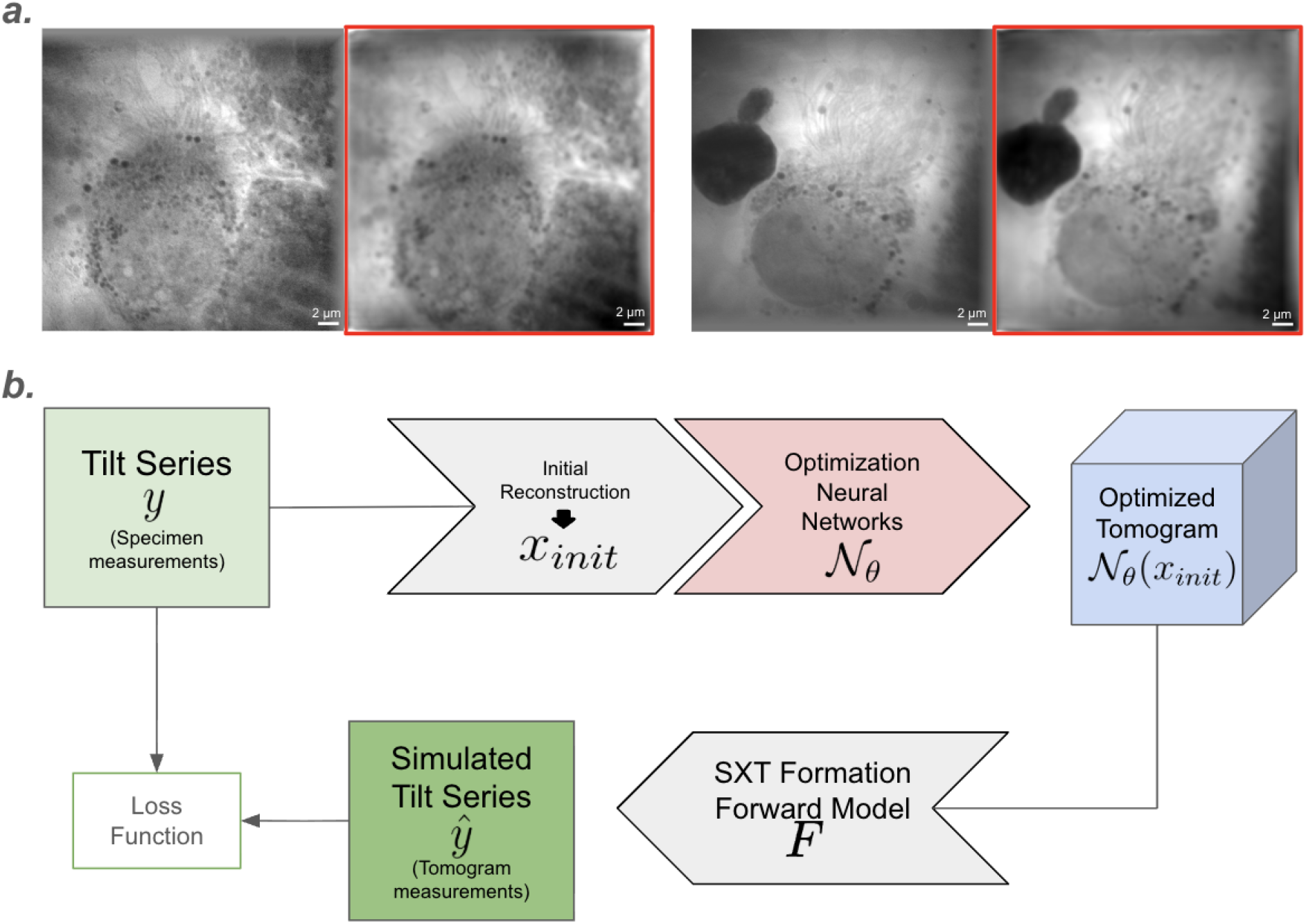
Degraded simulated tilt series (red boxes) derived with the SXT formation forward model and the measurement-supervised learning pipeline. a) Comparison of the original tilt series (measurements of the specimen) and simulated tilt series (red boxes, measurements of the reconstructed tomogram). The degradation of the simulated tilt series reflects the depth-dependent PSF mechanism modeled by the SXT forward model. b) Given the direct measurements of the specimen y, an initial reconstructed tomogram x_init_ is derived using conventional reconstruction algorithms (SIRT, WBP, etc.). x_init_ is then input to the optimization network N_θ_, parameterized by weights θ. Because the ground-truth biological specimen x does not exist in digital form, a direct loss between x and the network output cannot be computed. Instead, using the differentiable SXT forward model derived in equation (4), the loss is calculated in the measurement domain (tilt series), maintaining physical consistency with the optical degradation process throughout optimization.

Let *x* denote the true, unknown 3D biological specimen. The real experimental tilt series is acquired through the actual physical SXT imaging process, denoted as *F*_*real*_:

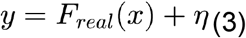

where *η* represents the measurement noise. The real tilt series *y* is first reconstructed into an initial 3D tomogram, *x*_*init*_, using conventional tomographic algorithms such as SIRT(Gilbert, 1972) or WBP(Radermacher, 1992). Due to the inherent optical limitations, *x*_*init*_ suffers from depth-dependent point spread function (PSF).

To recover the structure, we introduce a 3D neural network (e.g., 3D U-Net), denoted as *N*_*θ*_ and parameterized by its weights *θ*. The network aims to map the initial reconstruction to an optimized tomogram that approximates the real specimen:

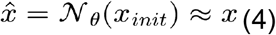

Different from standard Deep Image Prior (DIP) approaches, instead of initializing the network with random noise and training from scratch, the network takes an initial analytical reconstruction (*x*_*init*_), such as SIRT or WBP, as input. By optimizing directly on this structural baseline, the model bypasses the need to learn tomographic reconstruction physics and projection geometry. This restricts the network’s objective to correcting optical degradations and artifacts, which accelerates convergence and reduces overall inference time.

In an ideal supervised learning scenario, optimizing such a network requires minimizing the target distance between the network output and the ground truth:

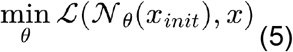

However, directly optimizing the neural network in the 3D spatial domain is not possible because the real biological specimen *x* does not exist in a digital ground-truth format.

To bypass the absence of ground-truth data, we shift the optimization from the 3D spatial domain to the 2D measurement domain. We utilize the differentiable forward model *F* derived in the previous section to computationally generate the simulated measurements *ŷ* of the network’s output:

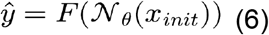

This allows us to optimize the neural network *N*_*θ*_ in a measurement-supervised manner. The optimal network weights 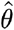 are obtained by minimizing the discrepancy between the computationally simulated measurements *ŷ* and the real experimental measurements *y*:

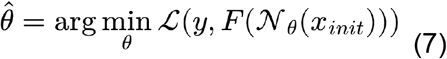

The rationale for this measurement-domain strategy is that *F* applies the known physical optical degradations to the network’s output 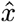 ; forcing the simulated degraded projection *ŷ* to match the real degraded projection *y* encourages the network to counteract those same degradations. If the network output 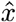 still contained residual defocus blur or artifacts inherited from *x*_*init*_, passing it through *F* would tend to produce a further-degraded projection that mismatches the real data more than a properly corrected volume would, favoring network outputs that compensate for the modeled PSF and recover high-frequency detail lost during imaging. However, because this inverse problem is ill-posed, and because both *N*_*θ*_ and *F* are necessarily imperfect approximations of the true imaging system, this optimization framework could still potentially introduce structured bias that aims to minimize the imperfect measurement loss. We address this concern empirically through split-tilt FRC analysis (in Results section) and, independently, through ground-truth validation using a bead phantom.

### Network Architecture

Processing full-resolution SXT tomograms (typically 1024*1024*1024 voxels) through a deep 3D neural network is computationally expensive and exceeds the 48 GB VRAM limit of the NVIDIA L40S GPU available on the Sonic HPC cluster at University College Dublin. To avoid the GPU memory bottleneck while preserving the original resolution in SXT, we designed a memory-efficient architecture that combines global residual learning(He et al., 2015) with a compact 3D U-Net backbone(Çiçek et al., 2016).

Specifically, the original dimension is drastically compressed from 1024^3^ to 256^3^ with 3D average pooling (scale factor of 4). This compressed volume is then processed by a compact 3D U-Net to compute a volumetric correction mask. Each convolutional block within the U-Net consists of a 3*3*3 convolution, 3D Batch Normalization, and an Exponential Linear Unit (ELU) activation. The feature channels start at 16 and double at each spatial downsampling step, reaching a bottleneck of 256 channels, before being decoded with skip connections. Most importantly, the computed correction mask is upsampled back to the 1024^3^ dimension and directly added to the original high-resolution input via a global skip connection. The key to preserving high-resolution details with sharp dimension compression during encoding lies in this residual design. Instead of forcing the network to learn the entire high-resolution volume from scratch, which would cause fine details to be lost inside the low-resolution bottleneck, the network only learns the necessary corrections. As a result, the original high-frequency features in the original input simply bypass the U-Net entirely and remain intact in the final output.

### Pretraining and Instance-Specific Fine-Tuning

A primary bottleneck of standard deep image prior (DIP)(Ulyanov et al., 2020) is the long training time required for the neural networks to learn tomographic reconstruction physics and projection geometry from scratch. To overcome this, our framework accelerates convergence by inputting an initial analytical reconstruction (e.g., SIRT or WBP) rather than random noise, which restricts the network’s objective strictly to correcting optical degradations rather than relearning projection geometry.

Building on this structural baseline, we propose a duo training-inference strategy. We first established a foundation model by pre-training the network on a diverse dataset of 97 SXT tomograms, using the measurement-supervised paradigm described above. During this stage, the model learns the general ultrastructures of diverse biological specimens. We adopt four different loss functions with different weights (L1 loss×1, SSIM loss×0.5(Zhou Wang et al., 2004), Fourier space loss×0.25(Fuoli et al., 2021), and gradient loss×1(Mathieu et al., 2015)) to ensure the preservation of physical features, enforcing both pixel-wise intensity consistency and high-frequency structural fidelity in the measurement domain. It is worth noting that the weights of different losses are currently empirical, selected based on preliminary experimentation rather than systematic optimization; a more rigorous exploration of loss selection and weighting, for example through ablation studies or automated weighting schemes, remains an important direction for future work.

To ensure physical fidelity of target tomogram in the inference stage, the framework is then fine-tuned specifically to an individual target tomogram (with the same losses and weights as described in the pre-training). In this stage, we adopt the DIP paradigm by further optimizing the network weights using the raw tilt series of the specific specimen at hand, following the measurement-supervised logic described in Equation (7).

Unlike the vanilla DIP, which may converge slowly or lack biological context, our approach utilizes both the initial analytical reconstruction as a structural input and the pre-trained weights as a starting point that incorporates structural knowledge learned from the 97-tomogram pretraining dataset. This pretrained prior is not removed by the subsequent fine-tuning stage; rather, fine-tuning further constrains it to be consistent with the specimen’s own direct measurements (tilt series). This is intended to support the recovery of high-frequency details that reflect the individual biological sample, though, as discussed above, the extent to which this constraint corrects rather than merely reshapes the pretrained prior cannot be fully confirmed by measurement-domain consistency alone, and is instead assessed empirically through the FRC and bead phantom validations presented in the Results.

### Computational Cost and Runtime

All computations were performed on the Sonic HPC cluster at University College Dublin using a single NVIDIA L40S GPU (48 GB VRAM). Pre-training the network on the 97-tomogram foundation dataset required approximately 15 hours for 20 epochs. While we empirically observe effective convergence during pre-training, whether additional or fewer epochs would yield better performance or efficiency remains an open question for future work. Instance-specific fine-tuning on an individual full-resolution tomogram, following the measurement-supervised optimization described above, required approximately 20 minutes per specimen (30 epochs), reaching a peak GPU memory usage of approximately 46 GB, near the 48 GB capacity of the available hardware. While additional fine-tuning epochs may yield further gains, we did not observe significant improvement beyond 30 epochs. Once fine-tuned, generating the final optimized tomogram via a single forward pass through the network required less than 1 minute. However, because the framework requires instance-specific fine-tuning for each new tomogram, the per-specimen computational cost is dominated by the fine-tuning stage rather than inference. Given that peak memory usage already approaches the limit of a 48 GB GPU, scaling this approach to larger tomograms or deploying it on hardware with less VRAM would likely require further memory optimization, a practical consideration alongside runtime when comparing this framework’s throughput to conventional reconstruction algorithms such as WBP or SIRT.

### Quantitative Feature Validation via FRC on Split Tilt-Series

To assess whether the recovered high-frequency details are consistent with the underlying biological specimen rather than reconstruction noise, we used a split-tilt-series FRC. The raw tilt series is split into two subsets based on even and odd indices, ensuring that both subsets capture the identical underlying biological architecture while their noise statistics remain independent. Because noise profiles are random, they fail to cross-correlate in the frequency domain, whereas a reproducible structural signal does not. FRC therefore provides evidence that recovered high-frequency content is consistent across independent noise realizations; it does not, on its own, exclude the possibility of correlated biases introduced by applying the same network architecture and optimization procedure to both subsets, nor does it directly quantify deconvolution or deblurring capability.

Specifically, these two subsets were reconstructed and optimized independently. Intensity normalization and a 2D Hanning window were applied before the Fourier transform to suppress spectral leakage caused by edge discontinuities. The cross-correlation across spatial frequencies was computed using an analytical arc-based algorithm. After mitigating localized noise fluctuations via a uniform moving average filter, the spatial resolution limit of each slice was defined as the inverse of the spatial frequency where the smoothed FRC curve intersected the fixed 0.25 threshold. This slice-by-slice quantification allows visualization of resolution changes throughout the volume.

Because FRC alone cannot fully exclude correlated artifacts or directly confirm that recovered features reflect true specimen structure, we additionally validated the framework using a bead phantom tomogram with known, ground-truth geometry (see Results). This independent validation, which does not rely on FRC or any correlation-based metric, provides direct confirmation that the optimization does not fabricate structural features, and that closely spaced objects can be genuinely resolved rather than merely appearing sharper.

### Biological sample preparation

#### Chlamydomonas, hBEC, and Huh-7 culture

The culture of *Chlamydomonas* and human bronchial epithelial cells (hBEC) has been described previously(Martin et al., 1975; De Ceuninck Van Capelle et al., 2025). Huh-7 cells were seeded into U-bottomed non-adherent 96-well plates (Sphera, Nunclon) at 10,000 cells per well in Dulbecco’s Modified Eagle’s Medium (DMEM) supplemented with 10% heat inactivated fetal calf serum and non-essential amino acids, and centrifuged for 5 mins at 200 x g. 72 hours post-seeding, the medium was supplemented with 1.5% DMSO for ten days to induce Huh-7 polarisation as previously described(Brown et al., 2025). Medium was exchanged every three days for maintenance. The spheroids were dissociated by incubation with 0.05% trypsin-EDTA (Gibco) and gentle mechanical shearing. The single cell suspension was then reseeded onto F1 Finder TEM grids (AgarScientific). After 24 hours, grids were counterstained with Hoechst 33342 and fixed with 4% Paraformaldehyde / 1.5% Glutaraldehyde for 1 hour before plunge freezing.

#### P. falciparum culture

Long-term *in vitro*-adapted *P. falciparum* clone 3D7 was maintained in culture as described previously(Lopez-Perez & Seidu, 2022). In brief, parasites were grown at 5% hematocrit in serum-free RPMI-1640 medium and maintained at 37°C under controlled atmospheric conditions (2% O_2_, 5% CO_2_, 93% N_2_). Synchronized late-stage infected erythrocytes were purified by magnetic columns(Lopez-Perez & Olsen, 2022) and adjusted to 18% parasitemia and 1% hematocrit. After 2 hours and 4 hours at 37°C in serum-free RPMI-1640 medium, infected erythrocytes were collected, washed twice in phosphate-buffered saline (PBS), and the pellet was diluted to 50% hematocrit with PBS before freezing.

For cryo-preservation for SXT imaging, cell samples were applied to 3 mm TEM grids, preliminarily treated by glow discharging (Ar atmosphere, 30 seconds, 10 mA), and vitrified immediately by plunge-freezing in Leica GP2 freezer (6s blotting, 80% sample chamber humidity).

### Soft X-Ray Tomography Imaging

Tomographic data were gathered with a lab-based SiriusXT SXT-100 microscope. To acquire the tomograms, soft X-rays were passed through flat specimens that had been cultivated on transmission electron microscopy (TEM) grids. These projections were recorded over a tilt range of roughly -55° to +55°, utilizing 1° intervals for each step. The overall time required to capture a complete tomogram ranged from one to two hours.

## Results

### Enhancement of cellular structures in Huh-7

To evaluate the efficacy of our measurement-supervised framework, we first examined a human hepatoma (Huh-7) cell tomogram in detail. This specimen was selected for its structurally complex cellular ultrastructure, which offers a demanding test of the framework’s ability to recover fine detail. Also, its comparatively large thickness (ca. 8.64 µm) further allows resolution recovery to be assessed across a wide range of focal depths, as reflected in the core-volume FRC analysis (Fig. 3c). Given this complexity, we first reconstructed the tomogram using two conventional baselines, Filtered Back-Projection (WBP) and Simultaneous Iterative Reconstruction Technique (SIRT), to examine how our optimization performs against distinct and complementary sources of degradation: WBP’s high-frequency noise and SIRT’s low-pass blurring. As shown in Fig. 3a, our measurement-supervised framework recovers high-frequency structural details regardless of the initial baseline, with optimized tomograms from both WBP and SIRT recovering consistent ultrastructural features.

**Figure 3.**
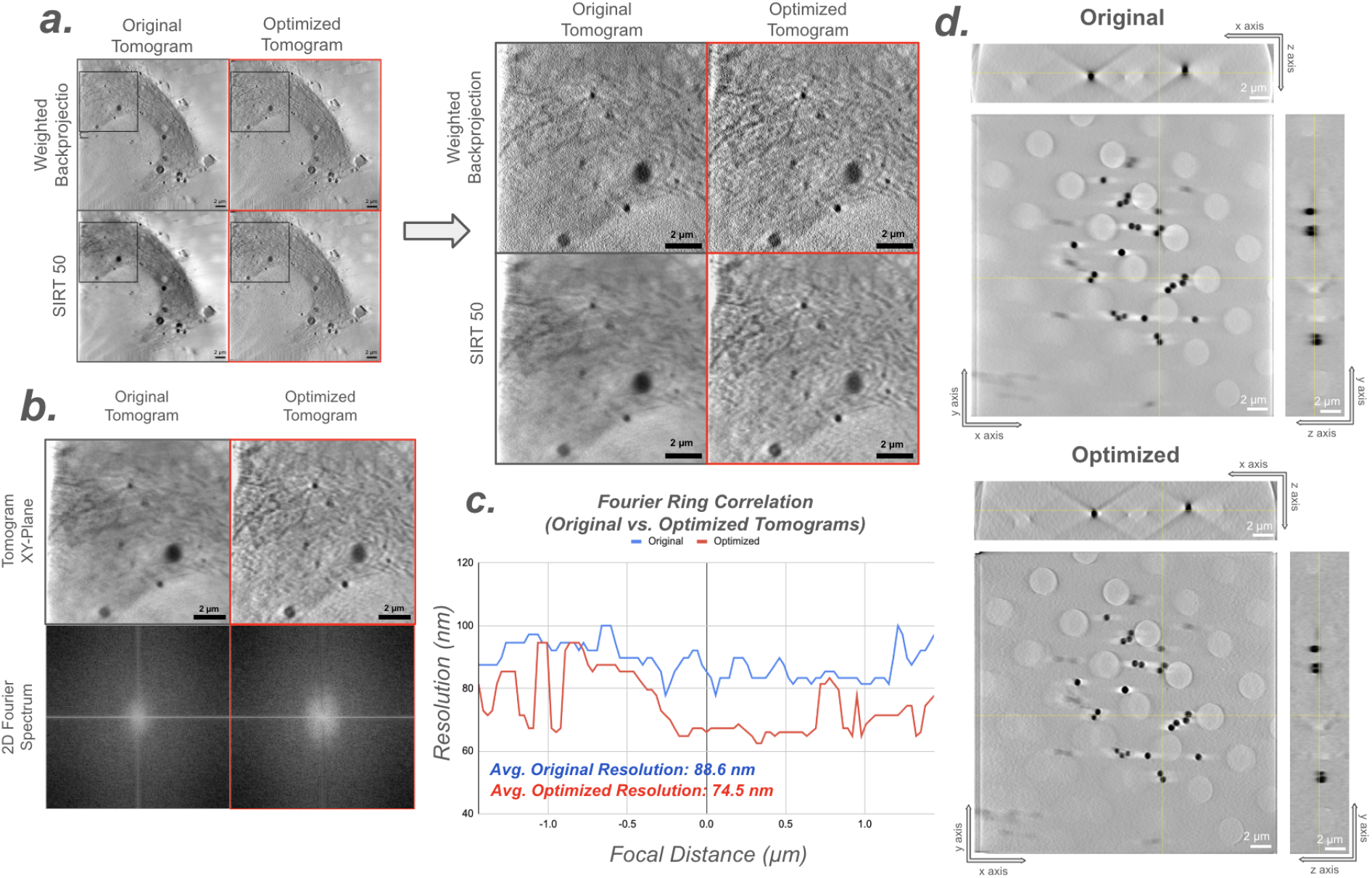
Examination of high-frequency ultrastructure recovery on Huh-7 and bead tomograms. a) Visual comparison of original baselines (WBP and SIRT) and our measurement-supervised optimization on a Huh-7 cell tomogram, demonstrating the recovery of high-frequency structural details regardless of the initial reconstruction baseline. b) 2D power spectra of localized regions comparing the SIRT baseline and the optimized tomogram. c) Fourier Ring Correlation (FRC) analysis on the Huh-7 tomogram, showing significant resolution improvements over the core cellular volume. d) Original and optimized reconstructions of isolated beads compared across three orthogonal planes (xy, xz, yz), showing that spherical shape is preserved after optimization.

To quantify this improvement, we computed the 2D power spectra of the localized regions using a fast Fourier transform (Fig. 3b). This frequency-domain visualization shows the expansion of high-frequency energy. We selected SIRT rather than WBP as the baseline for this and subsequent quantitative analyses because WBP reconstruction preserves high-frequency random noise that would hinder accurate FRC calculation. The 2D spectra illustrate that the SIRT baseline suffers from high-frequency attenuation caused by optical blurring, characterized by a confined central energy distribution. In contrast, our measurement-supervised optimization extends the spectral boundaries outward into the higher spatial frequencies.

However, an expanded spectral boundary alone does not guarantee the authenticity of these recovered features, as high-frequency energy could potentially stem from amplified noise or algorithmic artifacts. To verify that these recovered details correspond to genuine biological structures, we applied FRC to investigate the signal consistency, as detailed in the Methods section. By computing the FRC over the core cellular volume, we observed an average spatial resolution improvement from 88.6 nm to 74.5 nm in slices containing cellular structures (Fig. 3c). Because FRC quantifies signal reproducibility between two independently reconstructed and optimized sub-tomograms, this improvement indicates that the recovered high-frequency details are consistent rather than noise-driven, ensuring they reflect genuine biological structure. As detailed in the subsequent section (Fig. 4), this significant resolution improvement is consistently observed across an extensive dataset of nine tomograms featuring diverse biological ultrastructures.

**Figure 4.**
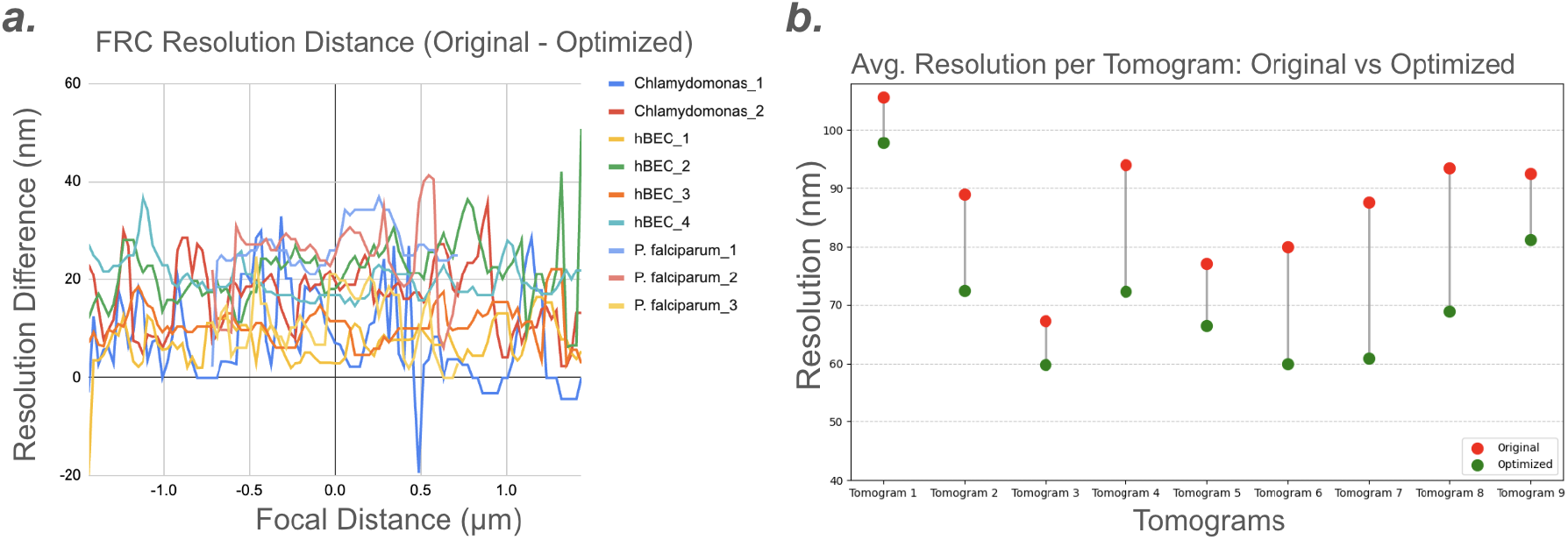
Quantitative FRC resolution improvements over 9 tomograms. a) Resolution difference across the core cellular volumes of nine independent tomograms. Positive values represent resolution gains (original − optimized), calculated via strictly independent split-tilt reconstructions and optimizations to ensure data integrity. b) Average absolute resolution gains: mean FRC resolution (in nm) before and after optimization for each of the nine tomograms, illustrating the magnitude of improvement relative to the varying baseline qualities of the individual samples.

While FRC confirms the recovery of reproducible biological signals, to further rule out the fabrication of geometric hallucinations, we evaluated the framework on a bead phantom tomogram providing a known ground truth. Comparing the original and optimized reconstructions across three orthogonal planes (xy, xz, and yz; Fig. 3d), the isolated beads retain their spherical geometry. We observed no evidence of anisotropic elongation or spurious internal substructures, confirming that the optimization restores existing features degraded by the forward model without introducing artificial shapes.

Because our measurement-supervised optimization framework can be grounded upon any conventional reconstruction algorithm, cross-benchmarking every existing method quantitatively is less central to our aims than quantifying the relative improvement over the specific baseline it is based upon. Therefore, for our quantitative analysis, we focused on how much our optimization improves compared to its initial baseline. We selected SIRT (50 iterations) as the optimization baseline, since SIRT acts as a low-pass filter that suppresses random high-frequency noise, helping to ensure accurate FRC calculations. Building on this baseline, our measurement-supervised framework mitigates residual point-spread-function blur by anchoring the optimization network to the specimen’s direct measurements (tilt series) through the differentiable SXT forward model.

### Generalization across complex biological ultrastructures

To systematically evaluate the generalizability of our framework, we applied the measurement-supervised optimization to another nine tomograms, encompassing Huh-7 cells, human bronchial epithelial cells (hBEC), Chlamydomonas, and P. falciparum. Due to varying sample thicknesses, the range containing cellular structures differs among the nine tomograms. To ensure the FRC quantification was performed on volumes containing meaningful cellular ultrastructure and to exclude empty background regions, we defined a specimen-specific core volume for each tomogram. The center of this volume was anchored to the optimal-focus slice, identified quantitatively as the slice yielding the highest spatial resolution (lowest value in nm). From this center, a symmetric volumetric window, typically spanning ±25 (0.72 μm) to ±50 slices (1.44 μm), was established.

We applied FRC analysis to the nine tomograms over their core cellular volumes (Fig. 4a). To prevent data leakage, the original tilt series of each specimen was split into two interleaved sub-stacks, which were independently reconstructed and optimized before FRC calculation. For visualization, we plotted the slice-wise resolution difference, calculated by subtracting the optimized resolution from the SIRT baseline at each Z-slice. As shown in Fig. 4a, the framework shows a significant improvement in spatial resolution across the core volumes of all nine specimens, with most difference values above 0. The correlated high-frequency signals between the independently optimized sub-tomograms further support that the recovered structures largely reflect reproducible signals rather than noise.

Because adjacent Z-slices within a tomogram are highly correlated and do not constitute independent replicates, we treated each tomogram, rather than each slice, as the unit of statistical replication. For each of the nine tomograms, we computed the mean FRC resolution across all slices in the core volume, separately for the SIRT baseline and the optimized reconstruction (Fig. 4b), and compared these nine paired values using a two-tailed paired t-test. This revealed a significant improvement in resolution following optimization (mean improvement = 16.3 nm, 95% CI: 10.7–21.9 nm; t(8) = 6.70, p = 0.00015, n = 9 independent tomograms; Cohen’s d = 2.23), consistent in direction across all nine specimens and robust to the exclusion of any individual tomogram. Given the modest sample size, we note that validation across a larger and more diverse specimen cohort would further strengthen confidence in the generality of this result.

To complement the quantitative FRC measurements, we qualitatively compared our framework against standard reconstruction baselines (Fig. 5). For this comparison, we selected Chlamydomonas and P. falciparum tomograms, as their densely packed and structurally complex cellular landscapes make differences between reconstruction methods more apparent than in less complex specimens.

**Figure 5.**
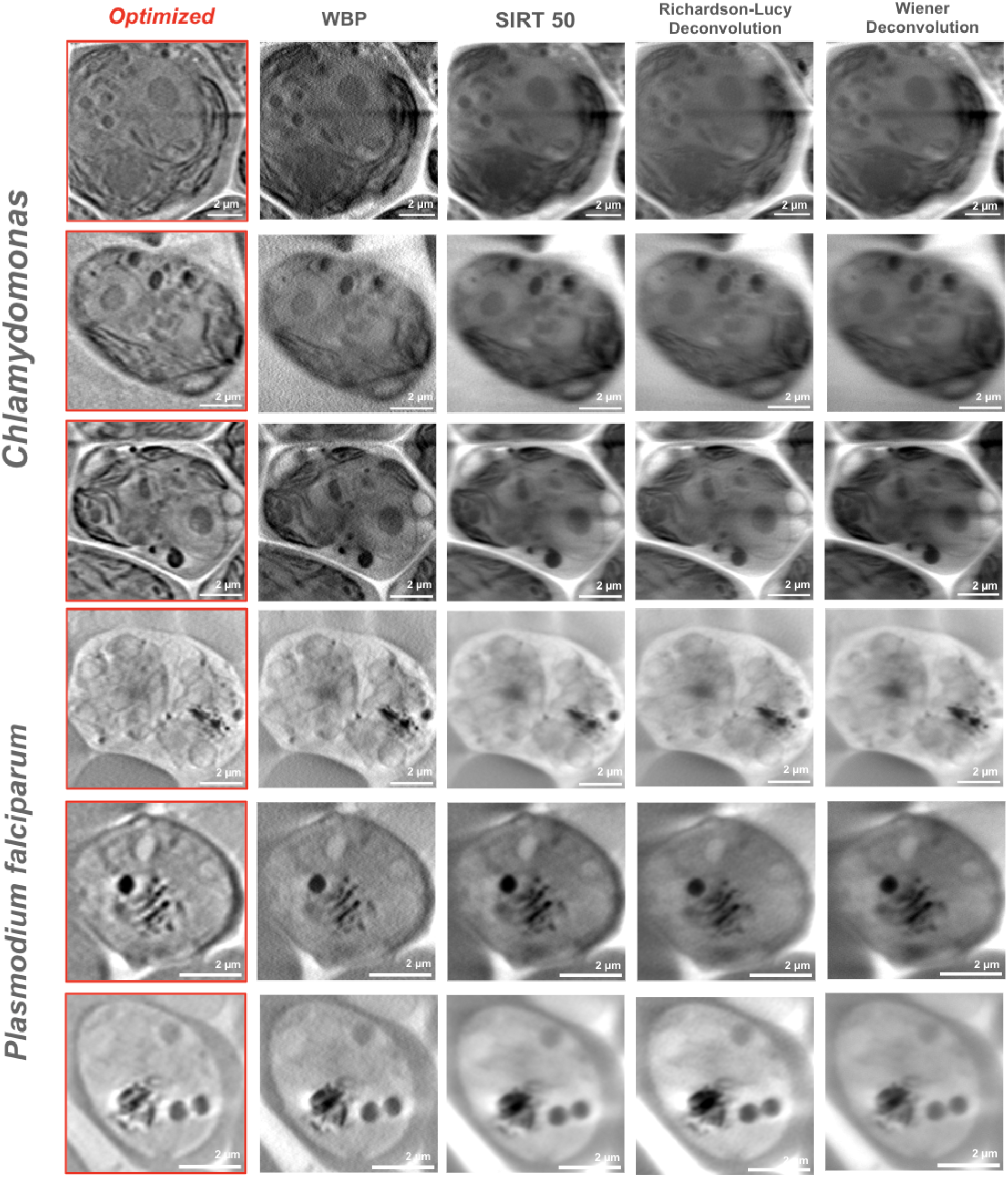
Qualitative comparison between standard SXT reconstructions and our measurement-supervised deep image prior (DIP) optimization. Representative tomograms from Chlamydomonas cells and Plasmodium falciparum demonstrate the framework’s performance on complicated cellular ultrastructures.

We first examined four conventional reconstruction approaches: WBP, SIRT (50 iterations), Richardson-Lucy, and Wiener deconvolution applied atop these baselines. WBP retains substantial high-frequency content, producing sharp cellular boundaries, but this comes with a correspondingly higher noise level that can obscure subtle ultrastructural detail. SIRT, in contrast, acts as a low-pass filter that effectively suppresses this noise but blurs high-frequency detail needed for accurate structural assessment. Applying Richardson-Lucy or Wiener deconvolution to a SIRT-50 reconstruction partially restores high-frequency signal, but the resulting structural clarity does not reach the level seen in WBP. Applying the same deconvolution methods directly to WBP did not yield obvious improvement in our experiments: WBP already retains most of the recoverable high-frequency content, leaving little additional structure for deconvolution to restore, while its existing noise is instead further amplified.

Our measurement-supervised optimization, optimized upon the SIRT-50 baseline, recovers a level of high-frequency structural detail comparable to WBP, while retaining SIRT’s suppressed noise floor (Fig. 5). We attribute this to the measurement-domain loss itself: because the network’s output is evaluated against the real tilt series via the differentiable forward model, rather than against the smoothed SIRT-50 reconstruction it was initialized from, the optimization is driven to recover high-frequency signal actually present in the raw measurements but suppressed by SIRT’s regularization, rather than being limited to whatever detail the initial reconstruction retained. This recovery is performed through tomogram-specific fine-tuning, meaning the network adapts directly to the measurements (tilt series) of each individual specimen rather than relying on a fixed, generically learned correction.

To directly examine this noise-suppression of our optimization against WBP, we compared line profiles drawn across major cellular structures in the Chlamydomonas and P. falciparum tomograms, reconstructed with WBP versus our optimization framework (Fig. 6). WBP line profiles show pronounced local intensity fluctuations that can obscure genuine underlying features, whereas the optimized tomogram yields smoother profiles with markedly reduced local fluctuation, consistent with effective noise suppression alongside preserved structural detail.

**Figure 6.**
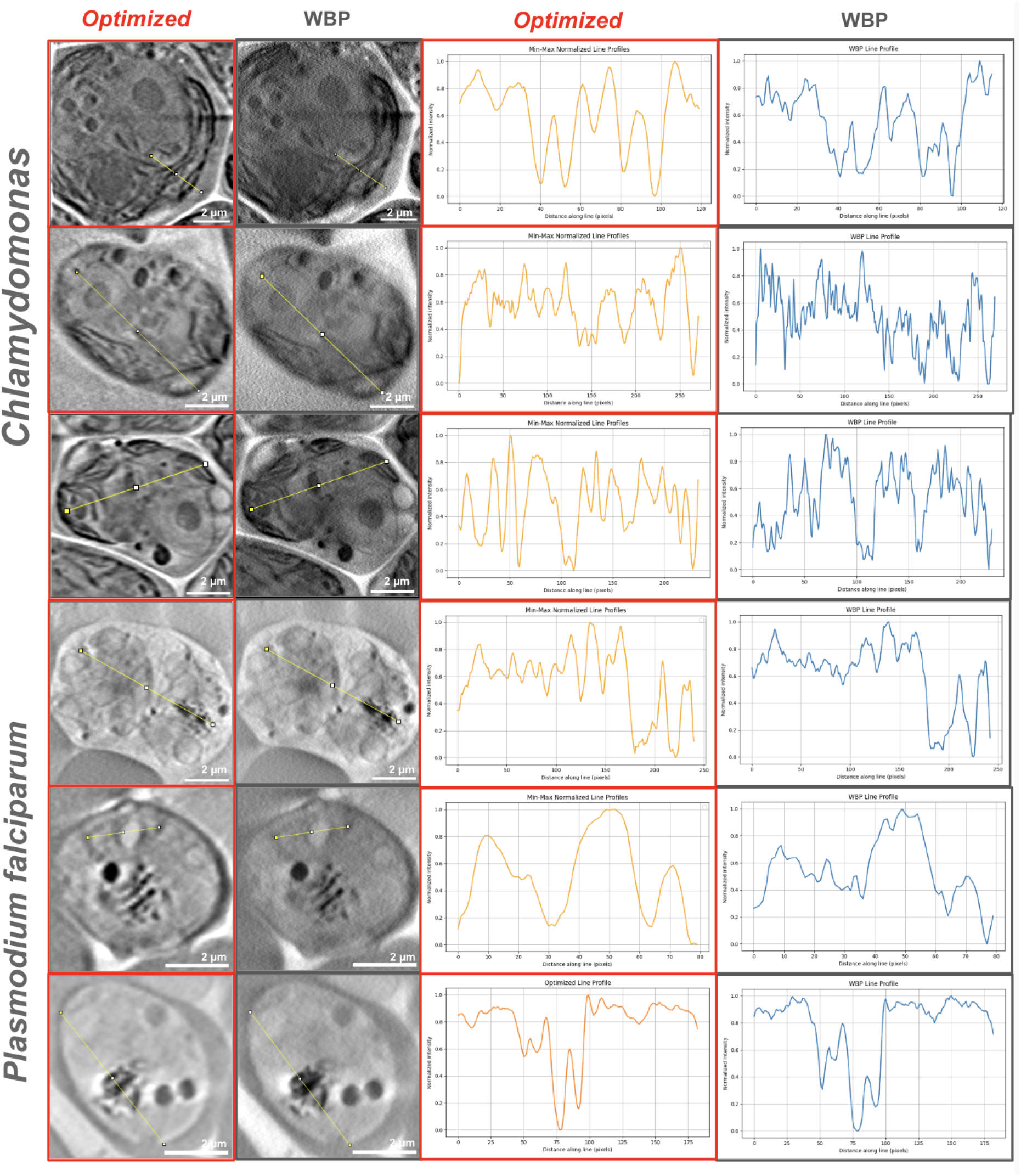
Line profile comparison between our optimized tomogram (SIRT-50 baseline) and WBP, drawn across major cellular structures in the Chlamydomonas and P. falciparum tomograms. WBP line profiles show pronounced local intensity fluctuations that can obscure genuine underlying features, whereas the optimized tomogram yields smoother profiles with markedly reduced local fluctuation, consistent with effective noise suppression alongside preserved structural detail.

### Demonstrating the potential for reduced-exposure imaging via sparse-angular subsampling

To explore the framework’s potential to support reduced-exposure imaging, we evaluated its performance under retrospective sparse-angular subsampling, in which fewer projection angles are drawn from an existing, full-exposure tilt series. In principle, the total radiation exposure delivered to a specimen can be reduced along two distinct axes: (i) reducing the photon flux delivered at each individual tilt angle while retaining the full angular sampling density, or (ii) retaining the full per-projection exposure while increasing the angular increment between projections, thereby reducing the total number of projections and the cumulative exposure integrated over the tilt series.

These two strategies are not interchangeable within our framework. Because the measurement-supervised loss (Eq. 7) is computed directly against the sampled tilt series, its effectiveness depends on each retained projection carrying sufficient structural clarity to provide an informative supervisory signal. Reducing the exposure per projection would degrade that signal directly and disproportionately undermine the measurement-domain optimization. Angular undersampling, by contrast, leaves the retained projections at full per-projection exposure and clarity while reducing only the number of viewing angles acquired.

Our retrospective subsampling experiment, which draws a reduced number of full-quality projections from an existing tilt series, is therefore best understood as a proxy specifically for angular undersampling (for example, acquiring every 2° rather than every 1°) rather than for exposure-per-angle reduction. This remains a retrospective analysis of already-acquired, full-exposure data and does not reproduce differences in sample motion, drift, or radiation-induced degradation that would accompany a genuinely shorter acquisition. However, because coarsening the angular increment is directly realizable without any hardware modification, this framing connects our retrospective analysis to a concrete, practical route for reducing radiation exposure in real acquisitions.

To assess resolution under a 2°-per-step angular sampling density (half the density of the original 1° tilt series), we followed the same split-half logic used for our full-angle FRC analysis, extended one level further. Estimating the resolution of a full, N-projection, 1°-step reconstruction requires splitting that series into two independent interleaved sub-stacks (each at an effective 2° spacing, N/2 projections), which are reconstructed and optimized independently and cross-correlated via FRC. To analogously estimate the resolution achievable from a combined N/2-projection, 2°-step tilt series, we split the original tilt series into four interleaved sub-stacks (each at an effective 4° spacing, N/4 projections), and randomly selected two of the four to serve as the independent split-halves for FRC calculation. Each selected sub-stack was reconstructed and optimized independently; their combined angular density (N/4 + N/4 = N/2 projections) is equivalent to that of a genuine 2°-step acquisition, and the FRC computed between them estimates the resolution attainable at that sampling density, following the same convention used in the full-angle case. We calculated the FRC resolution of these independently optimized sparse-angular tomograms and compared them to the standard full-angle SIRT baseline and the optimized full-angle reconstruction.

As Fig. 7a shows, the optimized sparse-angular tomograms match or exceed the slice resolution of the original full-angle baseline, generally falling between the standard full-angle baseline and the optimized full-angle upper bound.

**Figure 7.**
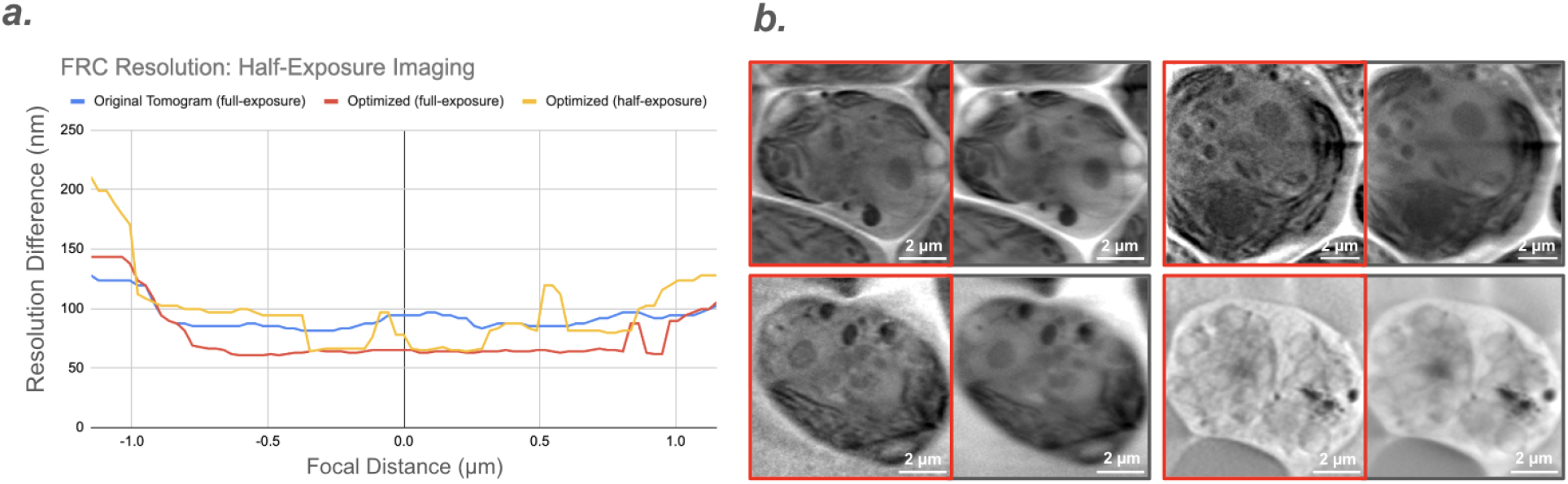
Performance under sparse-angular subsampling. (a) FRC resolution comparison among the standard full-angle baseline, optimized full-angle, and optimized sparse-angular tomograms (estimated via split-half FRC between two randomly selected quarter-density sub-stacks, together equivalent to a 2°-step acquisition), showing that the sparse-angular reconstruction matches the resolution of the full-angle baseline despite half the angular density. (b) Visual comparison between the optimized sparse-angular tomogram (red box) and the original full-angle tomogram. Although the initial, pre-optimization sparse-angular reconstruction is subject to streak artifacts characteristic of angular undersampling, the optimized reconstruction preserves fine cellular detail while suppressing these artifacts, consistent with the smooth spatial prior imposed by the network architecture.

We visualize the sparse-angular-optimized tomograms alongside the original full-angle tomogram to more intuitively assess visual fidelity. Despite using half the angular density, the recovered cellular structures remain comparable to those in the full-angle baseline. While these sparse-angular reconstructions show marginally higher noise than the optimized full-angle upper bound, high-frequency details are largely preserved.

This performance suggests the framework compensates for angular sparsity through the properties of the DIP. Standard analytical reconstructions typically suffer from pronounced streak artifacts when tilt angles are reduced, since the inverse problem is left with insufficient data constraints to unambiguously fill missing angular information. In our framework, the initial reconstruction supplied to the network is not exempt from this effect and can retain streak-like artifacts arising from the reduced angular coverage. However, because the 3D convolutional network architecture imposes an implicit, spatially smooth prior over the volume, consistent with prior DIP literature, the iterative, measurement-supervised optimization tends to suppress these unstructured streak patterns as it converges, favoring outputs consistent with both the network’s smooth structural prior and the retained, full-clarity projections.

Because each retained projection continues to provide the structural clarity needed for effective measurement-domain supervision, the network can leverage the available angular information to interpolate a resolution-consistent volume rather than propagating the sparse-angle artifacts present in the initial reconstruction. This dependence on per-projection clarity is also why we consider angular undersampling, rather than reduced exposure per angle, to be the acquisition strategy most directly supported by the present results. Confirming this potential for practical exposure reduction will require dedicated experiments using genuine coarse-angular acquisitions (e.g., 2° increments) with correspondingly reduced total scan-related sample degradation, which we identify as an important direction for future work.

## Discussion

The effective resolution of SXT is limited by depth-dependent optical degradation characterized by the PSF. By embedding this PSF mechanism into a differentiable forward model, our framework performs targeted PSF mitigation rather than generic image sharpening, allowing the recovery of high-frequency cellular ultrastructure that is otherwise lost to defocus blur. We emphasize that this framework does not eliminate the priors embedded in the network architecture, the 97-tomogram pretraining distribution, or the analytical initialization (SIRT or WBP); the recovered tomogram remains shaped by all of these choices, as it would in any learned or model-based reconstruction approach applied to an ill-posed inverse problem. What anchoring the optimization to raw measurements (tilt series), rather than to a fixed reconstructed volume, provides is a physically grounded consistency constraint: during instance-specific fine-tuning, the network’s output is required to remain compatible with the specimen’s own tilt series as passed through the measured forward model, rather than being free to drift arbitrarily far from it. This constraint narrows, but does not remove, the space of outputs consistent with the pretrained prior, and its practical effect on structural fidelity is something we assess empirically, rather than assume, through the split-tilt FRC and bead phantom validations described below.

This measurement-supervised strategy allowed the framework to consistently enhance resolution across nine independent tomograms with complicated cellular structures. Beyond the FRC-based quantification, our qualitative comparisons against classical baselines (WBP, SIRT, Richardson-Lucy, and Wiener deconvolution) helped intuitively observe the improvements. Post hoc deconvolution applied atop either WBP or SIRT only partially recovered high-frequency content and could not simultaneously match WBP’s sharpness and SIRT’s low noise floor. In contrast, our optimization recovered high-frequency detail comparable to WBP while retaining SIRT’s noise suppression, and the accompanying line-profile analysis showed reduced local intensity fluctuation relative to WBP. We attribute this to the measurement-domain loss itself: because the network is evaluated against the real tilt series rather than the smoothed reconstruction it was initialized from, it is driven to recover high-frequency signal present in the raw measurements but suppressed by the initial reconstruction’s regularization, rather than being limited to whatever a fixed deconvolution kernel can restore from an already-processed image.

To objectively verify the physical fidelity of these enhancements, we relied on two independent lines of evidence. First, split-tilt FRC from independently reconstructed and optimized sub-tomograms showed a statistically significant improvement across all nine specimens (mean improvement = 16.3 nm, 95% CI: 10.7–21.9 nm; t(8) = 6.70, p = 0.00015, n = 9 independent tomograms; Cohen’s d = 2.23), indicating that the recovered high-frequency content is reproducible across independent noise realizations rather than noise-driven. Second, because FRC alone cannot exclude correlated bias introduced by applying the same network and optimization procedure to both sub-tomograms, we additionally validated the framework using a bead phantom with known ground-truth geometry. The preserved spherical shape of isolated beads after optimization indicates the network does not fabricate structure where none exists. Together, these two validations, one based on signal reproducibility and one on known ground-truth geometry, provide converging, though not absolute, support that the recovered detail reflects genuine specimen structure.

Under retrospective sparse-angular subsampling, the framework maintained FRC resolution comparable to the full-angle baseline while using only half the projection angles. We interpret this as evidence that the 3D network architecture acts as an implicit spatial regularizer that helps compensate for missing angular information, consistent with prior observations in the DIP literature regarding resistance to streak artifacts. We acknowledge that this subsampling experiment is a computational proxy applied retrospectively to already-acquired, full-dose data, and does not reproduce the altered noise statistics, sample motion, or radiation-induced degradation that a genuinely reduced-exposure acquisition would introduce. To verify the practicability of true low-dose imaging requires further experiments with image acquisition pipeline, which we identify as an important direction for future work.

It is also worth noting that the forward model implementation in this study is based upon the experimentally measured PSF of the lab-based SXT microscope SXT-100, and directly applying the forward model to images acquired on other SXT microscopes will induce optical model mismatch, and could potentially lead to structural artifacts. While it is recommended that the PSF of each specific SXT microscope is experimentally measured to derive an accurate forward model, it is also possible to mathematically approximate the depth-dependent optical PSF(Oton et al., 2012).

## Conclusions

We demonstrated a physics-aware, measurement-supervised framework that recovers high-frequency cellular ultrastructure in soft X-ray tomography without modifying the imaging hardware or acquisition protocol. Across nine independent tomograms spanning four cell types, the framework produced a statistically significant improvement in spatial resolution, and this improvement was corroborated by two independent lines of evidence, split-tilt FRC and a bead phantom with known ground-truth geometry, supporting the conclusion that the recovered detail reflects genuine specimen structure rather than computational artifact. Under retrospective sparse-angular subsampling, the framework maintained resolution using half the projection angles, pointing to a practical, hardware-free route toward reduced radiation exposure in future acquisitions, though this remains to be confirmed with genuine low-dose acquisitions.

Our quantitative validation, while statistically significant, was based on nine tomograms acquired on a single lab-based instrument (SXT-100); extending this validation to a larger and more diverse specimen cohort, and to additional SXT systems, would further strengthen confidence in the generality and portability of the reported resolution gains. More broadly, the principles underlying this framework are not specific to SXT: PSF degradation and dose constraints affect other 3D imaging modalities, including cryogenic electron tomography (cryo-ET) and ptychographic X-ray computed tomography (PXCT), and coupling instrument-specific optical forward models with deep image priors offers a hardware-free approach that could help extend the spatial and dosimetric limits of nanoscale tomography more broadly.

## Author Contributions

***SSC***: Conceptualization, Data Curation, Formal Analysis, Investigation, Methodology, Software, Writing – Original Draft Preparation.

**EG, PS**: Methodology

***CdC, LL, TI***: Resources (biological samples), description of the biological sample preparation in the methodology section.

***CE, NF***: Resources (biological samples), description of the biological sample preparation in the methodology section.

***MLP***: Resources (biological samples), description of the biological sample preparation in the methodology section.

***CM, DR, SO, MD***: Conducting soft X-ray imaging of all samples presented in this work.

***JCS***: Supervision, Writing – Review & Editing

***SK***: Conceptualization, Formal Analysis, Validation, Supervision, Project Administration, Writing – Review & Editing.

## Data and Code availability

The data and code used in this study are available from Figshare (DOI:10.6084/m9.figshare.32408841, link: https://figshare.com/s/a1b83f425aff31c2faf6), and will be made fully publicly available upon acceptance.

Acknowledgments

SSC, JCS, and SK acknowledge funding from the European Union’s Horizon Europe Research and Innovation programme under the Marie Skłodowska-Curie Actions Doctoral Networks (MSCA-DN, agreement No. 101120151).

CdC, LL, and TI acknowledge funding from the Swiss National Science Foundation (IZLIZ3 200294).

## Declarations

The authors declare no competing interests.

## Notes

### Competing Interest Statement

The authors have declared no competing interest.

### Summary of Updates

Author Affiliation Change (last author affiliation: Copenhagen University => University College Dublin)

